# SCCA1/SERPINB3 promotes suppressive immune environment via STAT-dependent chemokine production, blunting the therapy-induced T cell responses

**DOI:** 10.1101/2023.02.01.526675

**Authors:** Liyun Chen, Victoria Shi, Songyan Wang, Rebecca Freeman, Fiona Ruiz, Kay Jayachandran, Jin Zhang, Pippa Cosper, Lulu Sun, Clifford J. Luke, Catherine Spina, Perry W. Grigsby, Julie K. Schwarz, Stephanie Markovina

## Abstract

Radiotherapy is a commonly used cancer treatment; however, patients with high serum squamous cell carcinoma antigen (SCCA1/SERPINB3) are associated with resistance and poor prognosis. Despite being a strong clinical biomarker, the modulation of SERPINB3 in tumor immunity is poorly understood. We investigated the microenvironment of SERPINB3 high tumors through RNAseq of primary cervix tumors and found that *SERPINB3* was positively correlated with *CXCL1/8, S100A8/A9* and myeloid cell infiltration. Induction of SERPINB3 *in vitro* resulted in increased CXCL1/8 and S100A8/A9 production, and supernatants from SERPINB3-expressing cultures attracted monocytes and MDSCs. In murine tumors, the orthologue *mSerpinB3a* promoted MDSC, TAM, and M2 macrophage infiltration contributing to an immunosuppressive phenotype, which was further augmented upon radiation. Radiation-enhanced T cell response was muted in SERPINB3 tumors, whereas Treg expansion was observed. A STAT-dependent mechanism was implicated, whereby inhibiting STAT signaling with ruxolitinib abrogated suppressive chemokine production. Patients with elevated pre-treatment serum SCCA and high pSTAT3 had increased intratumoral CD11b+ myeloid cell compared to patients with low SCCA and pSTAT3 cohort that had overall improved cancer specific survival after radiotherapy. These findings provide a preclinical rationale for targeting STAT signaling in tumors with high SERPINB3 to counteract the immunosuppressive microenvironment and improve response to radiation.

## Introduction

Radiotherapy (RT) is commonly used in the treatment of patients with squamous cell carcinomas, including the head and neck, esophageal, lung, and cervical cancer (1). RT can have both immunostimulatory and immunosuppressive effects, which in part determines the prognosis of cancer (1). The activation and infiltration of cytotoxic T cells post-radiation is critical to the curative activity of RT. However, tumors with an immunosuppressive tumor microenvironment (TME), dominated by myeloid cells tend to diminish T cell activity and may be more susceptible to the suppressive immune response induced by RT (2, 3). These immune responses include increased M2 macrophage polarization, myeloid-derived suppressor cell (MDSC) infiltration, T cell inhibitory receptor expression, as well as chemokine release upon radiation that alter the TME (4, 5). Chemokines are a subclass of cytokines with chemotactic properties that control the migration of cells and influence the composition of tumor immune microenvironment(6). Some chemokines promote an immunostimulatory environment, such as CXCL9, CXCL10, CXCL11, CXCL16, which improve dendritic cell activation and T cell trafficking to tumors (6, 7). Conversely, CCL2, CCL5, CXCL1, CXCL8, and CXCL12 can be induced by RT and have the opposite effect to recruit suppressive immune cells, inhibit effector T cells, and often correlate with poor treatment outcome (8-10).

Squamous cell carcinoma antigen 1 (SCCA), encoded by the *SERPINB3* gene locus and now known as SERPINB3, is a highly conserved cysteine proteinase inhibitor that interacts with lysosomal proteases upon lysosomal leakage and prevents cell death (11). We have recently demonstrated that SERPINB3 also protected cervix tumor cells against RT-induced cell death by preventing lysoptosis (12). In many cancers, SERPINB3/SCCA1 was highly expressed in tumors or in the circulation of cancer patients, including cervical, head and neck, lung, breast, and esophageal cancers, often associated with poor prognosis, treatment outcomes and recurrence (13-17). In addition, elevated SERPINB3 expression was also found in autoimmune disorders and implicated in the induction of inflammatory cytokines (18). However, in both tumors and autoimmune diseases the mechanistic link between SERPINB3 and immune regulation remains poorly understood. Considering the increasing number of studies reporting the association of SERPINB3 with tumorigenesis (19), metastasis (20), prognosis and recurrence, additional roles of SERPINB3, independent of proteinase-inhibitory activity, in tumor progression and resistance to therapy are likely.

We have previously demonstrated that patients with persistently high levels of SCCA before treatment and throughout the course of definitive RT had increased risk of recurrence and death (16). Prospective cohort studies also showed the prognostic value of SCCA for monitoring the response to RT and clinical outcome post-RT in cervical cancer patients (21). Given the unfavorable outcomes of patients with high SERPINB3 expression, we hypothesized that SERPINB3 promotes immune evasion by modulating suppressive immune responses that alter tumor microenvironment and impede RT-induced anti-tumor immunity. By characterizing cancer cells with high SERPINB3 expression within the context of the tumor microenvironment, we aim to provide a better understanding for treatment responsiveness and strategy to improve tumor control. Analysis of RNA-sequencing (RNAseq) data to predict immune infiltrate demonstrated enriched myeloid cell gene signature and immunosuppressive chemokine gene expression in tumors with high SERPINB3 expression. *In vitro* and *in vivo* assays suggested that chemokines induced by SERPINB3 contribute to myeloid cell migration and lead to a resistant environment to RT, in which high MDSCs, tumor-associated macrophages (TAMs), regulatory T cell (Treg) infiltration, and impaired T cell functionality were observed. We further discovered that SERPINB3 functioned through STAT signaling. Pharmacologic inhibition of STAT activity significantly reduced SERPINB3-associated suppressive chemokine production. Here, we present the clinical importance and regulatory function of SERPIN3 in establishing a pro-tumor microenvironment.

## Results

### SERPINB3 tumors are marked by myeloid cell-rich and suppressive immune profile

RNAseq was performed on 66 cervical tumor biopsies collected prior to (chemo)-RT. Patient and tumor characteristics of this cohort have been previously described, and are summarized in Supplemental Table 1. Patients were divided into three groups based on the distribution of SERPINB3 transcript levels; SERPINB3-low (B3/L, n=22), SERPINB3-intermediate (B3/Int, n=22), and SERPINB3-high (B3/H, n=22) groups (Figure 1A). To investigate the distinct immune signature associated with SERPINB3 expression in tumors, we focused our analysis on B3/L versus B3/H patient groups. The Immune Score (IS) was determined by xCell (22), via gene signature-based single-sample gene set enrichment analysis with the overall score representing a ranking of tumors in the dataset by lowest (IS of 0) to highest predicted immune infiltrate. B3/H tumors from patients who eventually experienced recurrence (R) compared to those who remained recurrence free (NR) showed overall higher immune scores compared to B3/L tumors indicating a potential immune-rich microenvironment (Figure 1B). Predicted immune cell content showed that B3/H tumors were characterized by increased myeloid cell subsets, including macrophages, monocytes, plasmacytoid dendritic cells and a small subset of CD8 T lymphocytes. In contrast, T-helper type 1 (Th1), Th2 and natural killer T cells were predicted to be lower in B3/H compared to B3/L tumors (Figure 1C).

**Figure 1.**
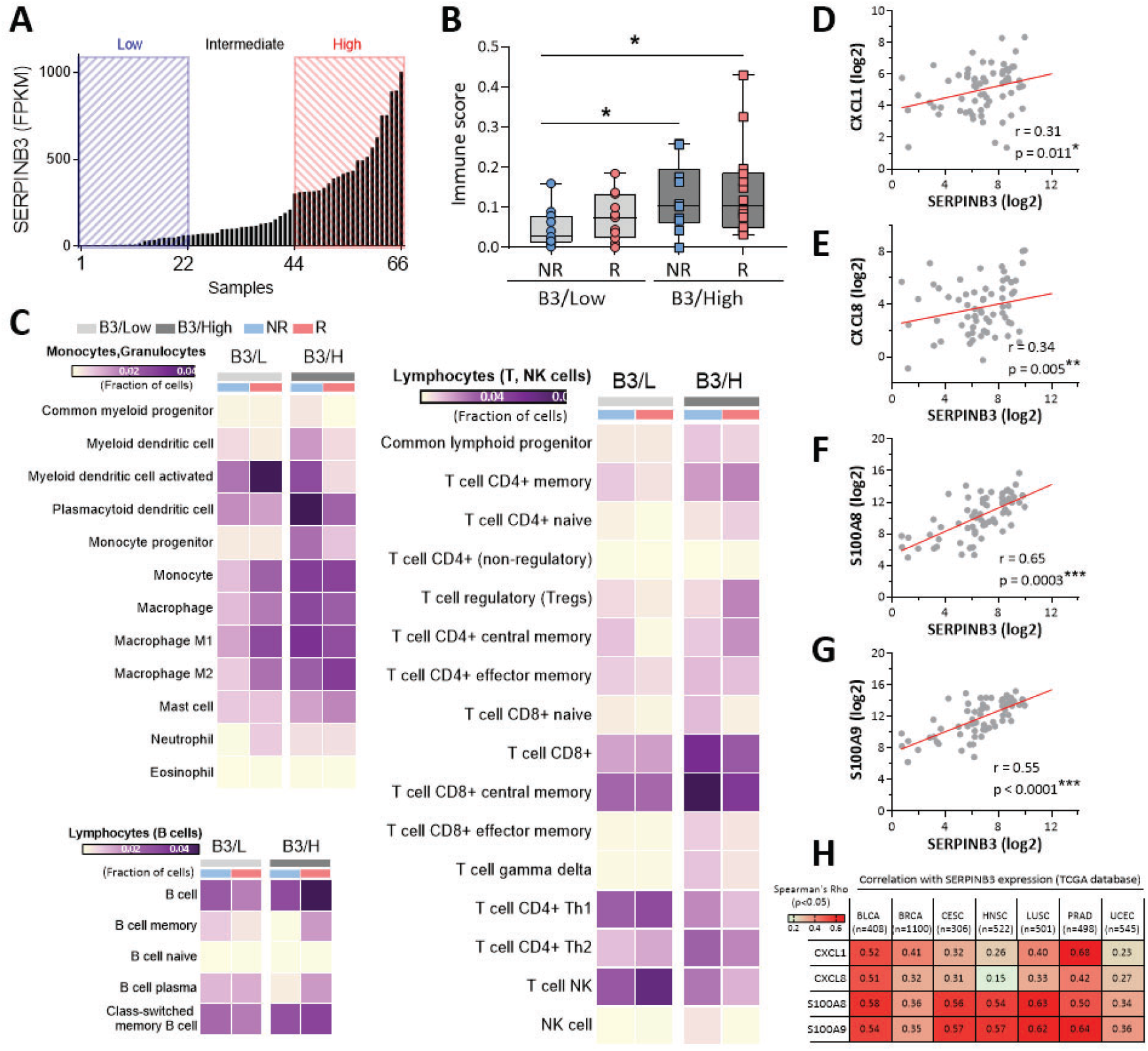
SERPINB3 tumors are marked by myeloid cell-rich and suppressive immune profile. (**A**) Normalized SERPINB3 transcript in cervical tumor biopsies from RNAseq was distributed by reads per kilobase of transcript per million mapped reads (RPKM). (**B**) Boxplots along with individual data points show xCell immune scores in recurrent (R)/non-recurrent (NR) SERPINB3-low (B3/L) and SERPINB3-high (B3/H) tumors. \**P* < 0.05, one-way ANOVA test. (**C**) Heatmap of enriched immune cell subpopulation was generated through xCell immune infiltrate prediction. Color intensity is proportional to average xCell score for each population across samples. (**D-G**) Spearman’s correlation of SERPINB3 with the expression of (**D**) CXCL1, (**E**) CXCL8, (**F**) S100A8, (**G**) S100A9 was performed using RNAseq from 66 cervical tumor biopsies collected prior to (chemo)- RT. (**H**) SERPINB3 expression correlated with CXCL1, CXCL8, S100A8, S100A9 expression in multiple cancer types. Analysis was performed using TCGA PanCancer Atlas and numeric values indicate Spearman’s correlation coefficient. BLCA, bladder urothelial carcinoma; BRCA, breast invasive carcinoma; CESC, cervical squamous cell carcinoma and endocervical adenocarcinoma; HNSC, head and neck squamous cell carcinoma; LUSC, lung squamous cell carcinoma; PRAD, prostate adenocarcinoma; UCEC, uterine corpus endometrial carcinoma

We then investigated differential expression of two major human chemokine subfamilies, CC and CXC chemokines, in the three groups (Supplemental Figure 1A). Two chemokines associated with myeloid cell migration - CXCL1 and CXCL8, correlated with SERPINB3 expression (Figure 1, D and E). In contrast, expression of T- and NK-cell recruiting chemokines CXCL9, 10 and 16, were not associated with SERPINB3 expression (Supplemental Figure 1A). Further analysis of chemokines that attract myeloid cells demonstrated a positive correlation between SERPINB3 and S100A8 / S100A9 expression (Figure 1, F and G). These correlations were validated in the TCGA-CESC (cervical squamous cell carcinoma and endocervical adenocarcinoma, n=306) dataset (Supplemental Figure 1B). Notably, analysis of the TCGA PanCancer Atlas showed a consistent positive correlation between SERPINB3 and myeloid-attracting chemokine expression across multiple tumor types including bladder, breast, head and neck, lung, prostate and uterine cancers (Figure 1H). These same tumor types have high levels of SERPINB3 expression (Supplemental Figure 1C).

### SERPINB3 results in upregulation of CXCL1/8 and S100A8/A9 chemoattractants, promoting myeloid cell migration from patient-derived peripheral blood

To study the mechanistic link between SERPINB3 and chemokine expression, we genetically altered SERPINB3 levels in human cervical cancer cells, Caski and SW756 cells (Supplemental Figure 2A), and examined the effect on chemokine production. Caski and SW756 with stable expression of SERPINB3 (Caski/B3, SW756/B3) showed increased CXCL1/8 and S100A8/A9 gene expression (Figure 2A) while downregulating SERPINB3 usingsh RNA (Caski/shB3) or CRISPR-Cas9-mediated deletion (SW756/CRISPR-B3KO) significantly reduced CXCL1/8 and S100A8/A9 expression (Figure 2B), when compared to their control counterparts. In addition to gene expression, significantly higher CXCL1/8 and S100A8/A9 protein expression and secretion was detected in Caski/B3 vs. Caski/Ctrl as well as SW756/B3 vs. SW756/Ctrl (Figure 2, C and D). Because Caski and SW756 are positive for HPV16 and HPV18 respectively, we examined whether SERPINB3-induced chemokine expression is associated with HPV infection using HPV-negative cervical cancer cells, C33A. Similar to the observation in HPV-positive cells, C33A with SERPINB3 upregulation (C33A/B3) showed increased CXCL1/8 and S100A8/A9 gene and protein expression, suggesting a HPV-independent mechanism (Supplemental Figure 2B). We next examined the chemotactic response of human peripheral blood mononuclear cells (PBMCs), obtained from seven patients with biopsy-proven cervical cancer prior to delivery of any treatment, to the chemokines secreted by tumor cells with high SERPINB3 expression using a transwell assay and flow cytometry to characterize migrated cells (Supplemental Figure 3A). Supernatant collected from Caski/B3 and SW756/B3 promoted the migration of CD11b^+^ myeloid cells, with an average of 1.9-fold increase in Caski/B3 vs. Caski/Ctrl and 2.1-fold increase in SW756/B3 vs. SW756/Ctrl, whereas the migration of CD4^+^ and CD8^+^ T cells showed no statistical difference (Figure 2, E and F). Further analysis of migrated CD11b^+^ cells showed that populations migrating in response to both Caski/B3 and SW756/B3 supernatant were enriched in monocytes, and monocytic and polymorphonucler myeloid-derived suppressor cells (M-/PMN-MDSCs) with an approximately 1.5-2 fold increase compared to Ctrl supernatant (Figure 2G and Supplemental Figure 3, B and C).

**Figure 2.**
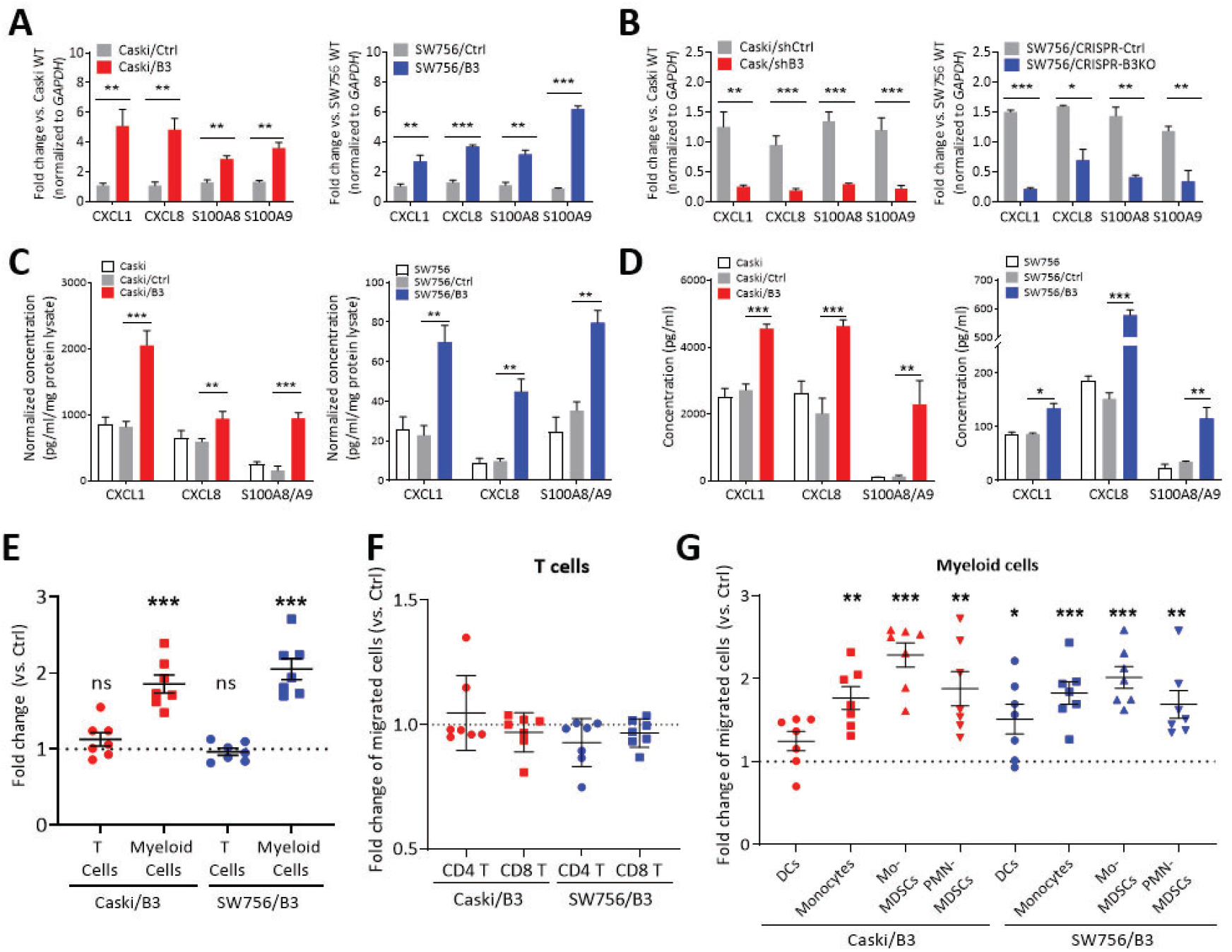
SERPINB3 results in upregulation of CXCL1/8 and S100A8/A9 chemoattractants, promoting myeloid cell migration from patient-derived peripheral blood. (**A**) Caski and SW756 cells were transduced with pUltra vector (Caski/Ctrl, SW756/Ctrl) or pUltra-SERPINB3 (Caski/B3, SW756/B3) and CXCL1/8 and S100A8/A9 mRNA expression was examined by qPCR. (**B**) Caski cells were transfected with scrambled negative control shRNA (Caski/shCtrl) or shRNAs specific SERPINB3 (Caski/shB3); SW756 cells were transduced with CRISPR control vector (SW756/CRISPR-Ctrl) or CRISPR-Cas9 for SERPINB3 knockdown (SW756/CRISPR-B3KO). The expression of CXCL1/8 and S100A8/A9 was examined by qPCR. Gene expression were normalized to GAPDH and fold changes were calculated by comparing to the expression levels in parental cells (Caski WT or SW756 WT). (**C**) Intracellular chemokine proteinexpressionin cell lysateswas measured by ELISA. The chemokine levels were normalized to total protein concentration. (**D**) Supernatant was collected from adherent cells in monolayer and chemokine secretion was measured by ELISA. Data in **A-D** are presentedas mean ± SEMof n = 4 independent experiments, \**P* < 0.05, \*\**P* < 0.01, \*\*\**P* < 0.001 using unpaired two-tailed Student’s t-test. (**E-G**) PBMC migration towards supernatant collected from cancer cells was examined by Transwell assays and the migrated PMBC populations were analyzed by flow cytometry. Fold changes were calculated as the percentage of migrated (**E**) T and myeloid cells, (**F**) T cell subsets and (**G**) myeloid cell subsets in Caski/B3 or SW756/B3 relative to Caski/Ctrl or SW756/Ctrl supernatant. Data are shown as mean ± SEM, ns, no significance; \**P* < 0.05, \*\**P* < 0.01, \*\*\**P* < 0.001 using two-tailed one sample T test against 1. Each dot represents the mean of duplicate values for a single donor sample (n=7).

### SERPINB3 tumors show accumulated myeloid cells and increased tumor growth

Since SERPINB3 upregulated the expression of myeloid chemoattractant *in vitro*, we hypothesized that tumor expressing SERPINB3 attract myeloid cell infiltration and mediate *in vivo* TME. We first injected human Caski/Ctrl or Caski/B3 cells subcutaneously on the flank of athymic nude mice, which do not have mature T cell compartments but accommodate human tumor cell growth and have functional myeloid cells. Tumor growth over time was quantified and tumor-infiltrating immune cells were analyzed using flow cytometry (Supplemental Figure 4A). Tumor growth showed no difference between Caski/Ctrl and Caski/B3 over the course of the experiment; however, Caski/B3 tumors had a significant increase in infiltrating CD11b+ myeloid cells compared to Caski/Ctrl tumors at days 22 and 40 post-injection (Supplemental Figure 4, B and C). M-MDSCs, TAMs, and M2 macrophages were significantly increased in Caski/B3 at both days 22 and 40, while no difference was seen in DCs, PMN-MDSCs and B cells (Supplemental Figure 4D).

Given that lymphocyte-mediated immune activities play a role in tumor response to radiotherapy, and that RT is known to reshape the TME, we next characterized SERPINB3-mediated TME and its response to radiation in an immunocompetent murine model. However, there are no murine cervical tumor lines, and the commonly used alternative, TC1 cells with HPV E6/E7 gene expression, were derived from normal lung epithelial cells with relatively low chemokine expression (Supplemental Figure 5A). Therefore, constructs driving murine *Serpinb3a*, homologous to human SERPINB3 (23), were expressed in LL2 murine lung carcinoma cells (LL2/mB3a) and an empty vector was used as a control (LL2/Ctrl) (Supplemental Figure 5B). Of note, SERPINB3 is also expressed in lung cancer (supplemental Figure 1C) and negatively associated with prognosis, providing credence to this model (13). As in the human cervical cancer model, the gene expression and secretion of murine Cxcl1/3, functionally corresponding to human CXCL1/8, and murine S100a8/9, homologous to human S100A8/9, were induced by mSerpinb3a, whereas chemokines associated with T cell migration, mCxcl9 and mCxcl10 were not affected by mSerpinb3a expression (Figure 3, A and B).

**Figure 3.**
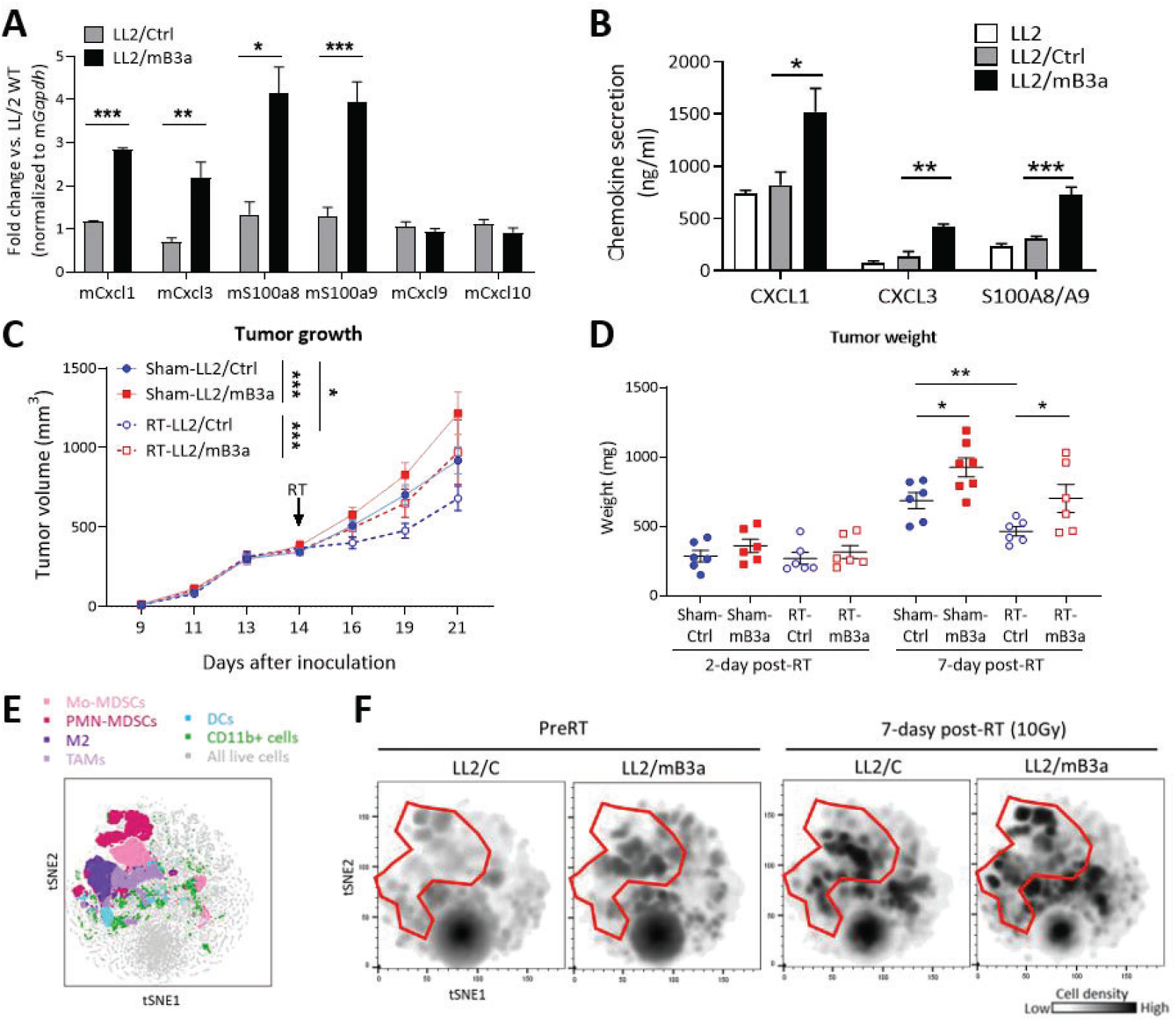
SERPINB3 tumors show accumulated myeloid cells and increased tumor growth. (**A**) Chemokine mRNA expression in LL2 cells transduced with pLV-C-GFPSpark vector with mSerpinb3a expression (LL2/mB3a) or a control pLV-C-GFPSpark vector (LL2/Ctrl) were examined by qPCR. Gene expression was normalized to mGapdh and fold changes were calculated by comparing to the expression levels in LL2 parental cells. (**B**) Chemokine secretion was measured by ELISA. Supernatant was collected from the adherent cells in monolayer at 72 hours. Data in **A-B** are shown as mean ± SEMof n = 4 independent experiments, \**P* < 0.05, \*\**P* < 0.01, \*\*\**P* < 0.001 using unpaired two-tailed Student’s t-test. (**C**) Tumor growth of C57/BL6 mice with LL2/mb3a tumors (red lines) and LL2/Ctrl tumors (blue lines) were randomized to receive sham-treated (solid lines) or 10Gy RT at day 14 (dotted lines). Significance was calculated using two-way ANOVA, \**P* < 0.05; \*\*\**P* < 0.001 (**D**) Tumor weight at the experimental endpoint were measured. The graph represents mean ± SEM with individual data points, n=6-7 mice in each group, \**P* < 0.05, \*\**P* < 0.01 using one-way ANOVA. (**E**) viSNE plots show flow cytometry analysis of total viable cells from tumors with separate clustering by pre-definedcell surface markers for myeloid cell subsets, including M-MDSCs (CD11b+Ly6G-Ly6Chigh), PMN-MDSCs (CD11b+Ly6G+), TAM (CD11b+Ly6G-F4/80+), M2 macrophages (CD11b+Ly6G-F4/80+CD163+) and DCs (CD11b+Ly6G-F4/80-CD11c+). viSNE plots were generated by concatenating individual flowcytometry data in each group into single file. The red-circled area indicates CD11b+ cells and expression levelsare shown by color in relative intensity.

LL2/Ctrl and LL2/mB3a cells were injected subcutaneously into C57/BL6 mice and mice with tumors were randomized to receive a single dose of 10 Gy or sham RT (14 days post-injection). Mice were sacrificed 2 days post-RT (day 16) and 7 days post-RT (day 21), with tumors and tumor-draining lymph nodes harvested. Although this experiment was not designed to detect tumor growth delay following radiation, RT-treated LL2/Ctrl tumors had the smallest tumor volumes, and RT-treated LL2/mB3a tumor volume curves overlapped with the sham-treated LL2/Ctrl tumor volume curve (Figure 3C). This is consistent with our prior study showing that human cervical tumor cell lines expressing SERPINB3 are more radioresistant than control tumors in an athymic nude murine model (12). Tumor weights showed no statistical differences in all groups at 2-day post-RT while a more substantial increase in sham- and RT-LL2/mB3a tumor growth was corresponded to increased tumor weight at 7-day post-RT compared to LL2/Ctrl counterpart. Radiation suppressed LL2/Ctrl tumor growth, reflected by reduced tumor weight at 7-day post-RT but RT-vs sham-LL2/mB3a was not statistically different (Figure 3D). Infiltrating immune cells were analyzed using flow cytometry (Supplemental Figure 6A) and an unsupervised analysis using the visualization of t-distributed stochastic neighbor embedding (viSNE) was exploited to cluster CD11b^+^ myeloid cell subsets based on pre-defined markers (Figure 3E). Prior to RT, CD11b^+^ myeloid cell density was higher in LL2/mB3a tumors compared to LL2/Ctrl. Irradiated tumors of both types had higher total CD11b^+^ myeloid cells, but different subsets were represented in LL2/Ctrl and LL2/mB3a (Figure 3F, red outline). Thus, we sought to examine further the dynamic change of myeloid cell subsets in LL2/Ctrl and LL2/mB3a tumors at different time points.

### SERPINB3 tumors are enriched for suppressive myeloid and Treg cells which is further augmented by radiation

Similar to *in vitro* findings, LL2/mB3a tumors had higher levels of intra-tumoral CXCL1 and S100A8/A9 expression over time, compared to Sham-LL2/Ctrl (Figure 4, A and B, solid bars). Radiation promoted further CXCL1 production in RT-LL2/mB3a but not in RT-LL2/Ctrl (Figure 4A). Although radiation induced S100A8/A9 in both RT-LL2/Ctrl and RT-LL2/mB3a tumors at 2 days post-RT, the magnitude of chemokine induction was greater in RT-LL2/mB3a than RT-LL2/Ctrl, with an average of 2.3-fold and 1.8-fold increase, respectively (Figure 4B). Higher and more persistent expression of immunosuppressive chemokines in the tumor milieu of irradiated LL2/mB3a tumors led us to hypothesize that the increased myeloid compartment summarized by viSNE plots differed specifically in immunosuppressive myeloid cell subtypes. Indeed, Sham-treated LL2/mB3a tumors showed consistently higher infiltration of M-MDSCs and PMN-MDSCs compared to LL2/Ctrl while radiation induced an early increase of infiltrating M- and PMN-MDSCs at 2-day post-RT in both groups; however, MDSCs in irradiated tumors remained elevated compared to sham-treated tumors at 7-days post-RT only in RT-LL2/mB3a tumors (Figure 4, C and D). The number of infiltrating TAMs and M2 macrophages was higher in sham LL2/mB3a versus LL2/Ctrl, with a gradual increase in both groups as the tumors grew, but no statistical change with irradiation in either genetic background (Figure 4, E and F). The ratio of M2 macrophages to total TAMs, however, was significantly higher in sham-treated LL2/mB3a compared to LL2/Ctrl, with a further increase induced by radiation only in the LL2/mB3a tumors (Supplemental Figure 6B). Tumor infiltrating DCs were not different between LL2/mB3a and LL2/Ctrl tumors. Although RT has been shown to enhance DC migration (3), a radiation-associated increase was seen only in RT-LL2/Ctrl tumors at 7 days post-RT, and not in LL2/mB3a (Supplemental Figure 6C).

**Figure 4.**
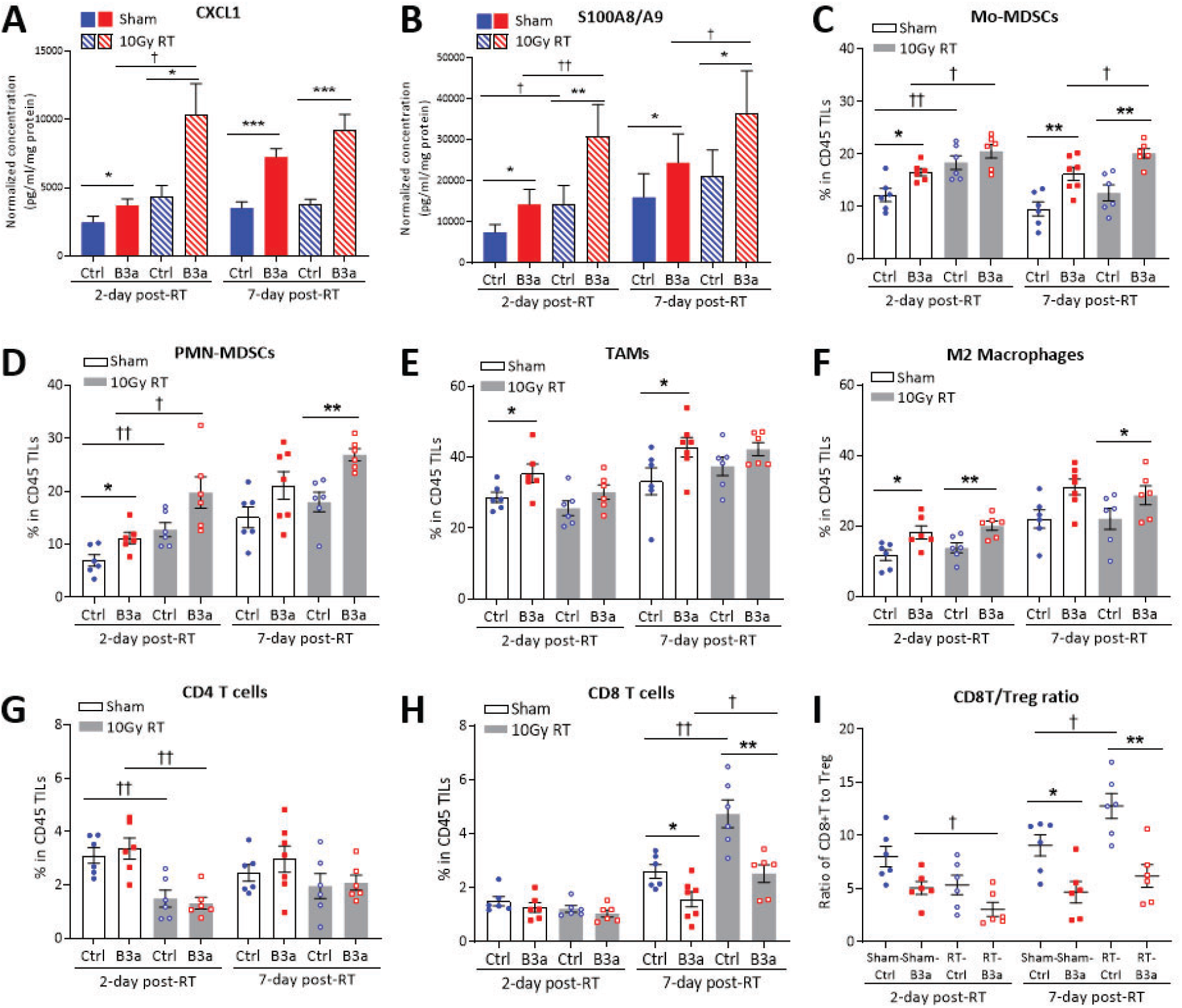
SERPINB3 tumors are enriched for suppressive myeloid and Treg cells which is further augmented by radiation. (**A-B**) Chemokine (**A**) CXCL1 and (**B**) S100A8/A9 levels in tumor homogenates was examined by ELISA. Data was normalized to the protein concentration for each tumor homogenate and shown as mean ± SEM; n=7 for 7-day sham-LL2/mB3a and n=6 for all other groups. (**C-H**) Cumulative data from FACS analysis show alteration of immune cell infiltration by SERPINB3 expression and radiation in LL2 tumors. The graphs represent the frequencies of (**C**) CD11b+Ly6G-Ly6Chigh Mo-MDSCs, (**D**) CD11b+Ly6G+ PMN-MDSCs, (**E**) CD11b+Ly6G-F4/80+ TAMs, (**F**) CD11b+Ly6G-F4/80+CD163+ M2 macrophages, (**G**) CD3+CD4+ T cells, and (**H**) CD3+CD8+ T cells in total tumor infiltrating leukocytes (TILs). (**I**) The ratioof CD8/Treg represented the infiltrating percentage of CD8+ T cells relative to CD4+CD25+FoxP3+ regulatory T (Treg) cells. Data are shown as mean ± SEM and each dot represents a biologically independent animal; * indicates comparisons between LL2/Ctrl and LL2/mB3a; ✝ indicates comparisons between sham-treated and RT; \**P* < 0.05, \*\**P* < 0.01, \*\*\**P* < 0.001 using one-way ANOVA with Tukey’s post-test for multiple comparisons.

The immunosuppressive environment in LL2/mB3a tumors might favor myeloid cell infiltration and prevent T cell recruitment. However, no difference was seen in CD4^+^ TILs between LL2/Ctrl and LL2/mB3a tumors and a significant decrease at 2-day post-RT in both groups was associated with radiation effect (Figure 4G), consistent with radiosensitivity of in-field lymphocytes (24). CD8^+^ TILs showed a comparable level at 2-day post-RT in all groups but significantly lower in Sham-/RT-LL2/mB3a vs in LL2/Ctrl at 7-day post-RT. Moreover, CD8^+^ TILs doubled in RT-LL2/Ctrl tumors compared to sham-treated tumors, and while statistically increased, the magnitude of increase was less in RT-LL2/mB3a (Figure 4H). Although the ratios of total CD8^+^ TILs were comparable in sham-/RT-LL2/Ctrl and LL2/mB3a tumors at 2-day post-RT, we found that the ratio of CD8^+^ T to Treg (CD4^+^CD25^+^FoxP3^+^) was significantly decreased in RT-LL2/mB3a compared to Sham-LL2/mB3a, indicating an increase of Treg cells in LL2/mB3a tumors shortly after radiation. In contrast, increased CD8^+^ TILs in RT-LL2/Ctrl at 7-day post-RT correlated with higher CD8^+^ T/Treg ratio compared to sham-LL2/Ctrl tumors (Figure 4I and Supplemental Figure 6D). Collectively, these data showed that the release of CXCL1 and S100A8/A9 was amplified upon radiation, perhaps by both SERPINB3-expressing tumor cells and increased suppressive myeloid cell infiltrates, while radiation-associated CD8^+^ T cell infiltration in SERPINB3 tumors was counteracted by an expansion of Treg cells, suggesting an exacerbated RT-induced immunosuppressive response.

### Tumor infiltrating T cells from mSerpinb3a tumors display impaired proliferation and functionally exhausted phenotype

With evidence of immunosuppression demonstrated by high infiltrating MDSCs, M2/TAM and Treg cells, T cell function is likely to be compromised in *mSerpinb3a* tumors. Moreover, as radiation is thought to enhance T cell responses (9), we also examined LL2/Ctrl and LL2/mB3a tumor-derived T cell proliferation and functionality following irradiation. The proliferation marker Ki-67 expression in CD8^+^ and CD4^+^ T cells was evaluated by flow cytometry (Supplementary Figure 7). At 2-day post-RT, despite comparable levels of CD8^+^ and CD4^+^ TILs, percent of Ki-67^+^ CD8^+^/CD4^+^ TILs were significantly lower in both Sham- and RT-treated LL2/mB3a compared to LL2/Ctrl tumors, indicating low proliferative capability (Figure 5, A and B). Tumor-directed radiation promoted CD8^+^ T cell infiltration but not proliferation at 7-day post-RT as most CD8^+^ T cells showed low Ki-67 expression in both LL2/Ctrl and LL2/mB3a tumors (Figure 5A). In contrast, CD4^+^ TILs that were initially decreased at 2-day post-RT in both groups rebounded slightly 7-day post-RT, and with enhanced Ki-67 expression in both RT-LL2/mB3a and RT-LL2/Ctrl compared to sham groups (Figure 5B).

**Figure 5.**
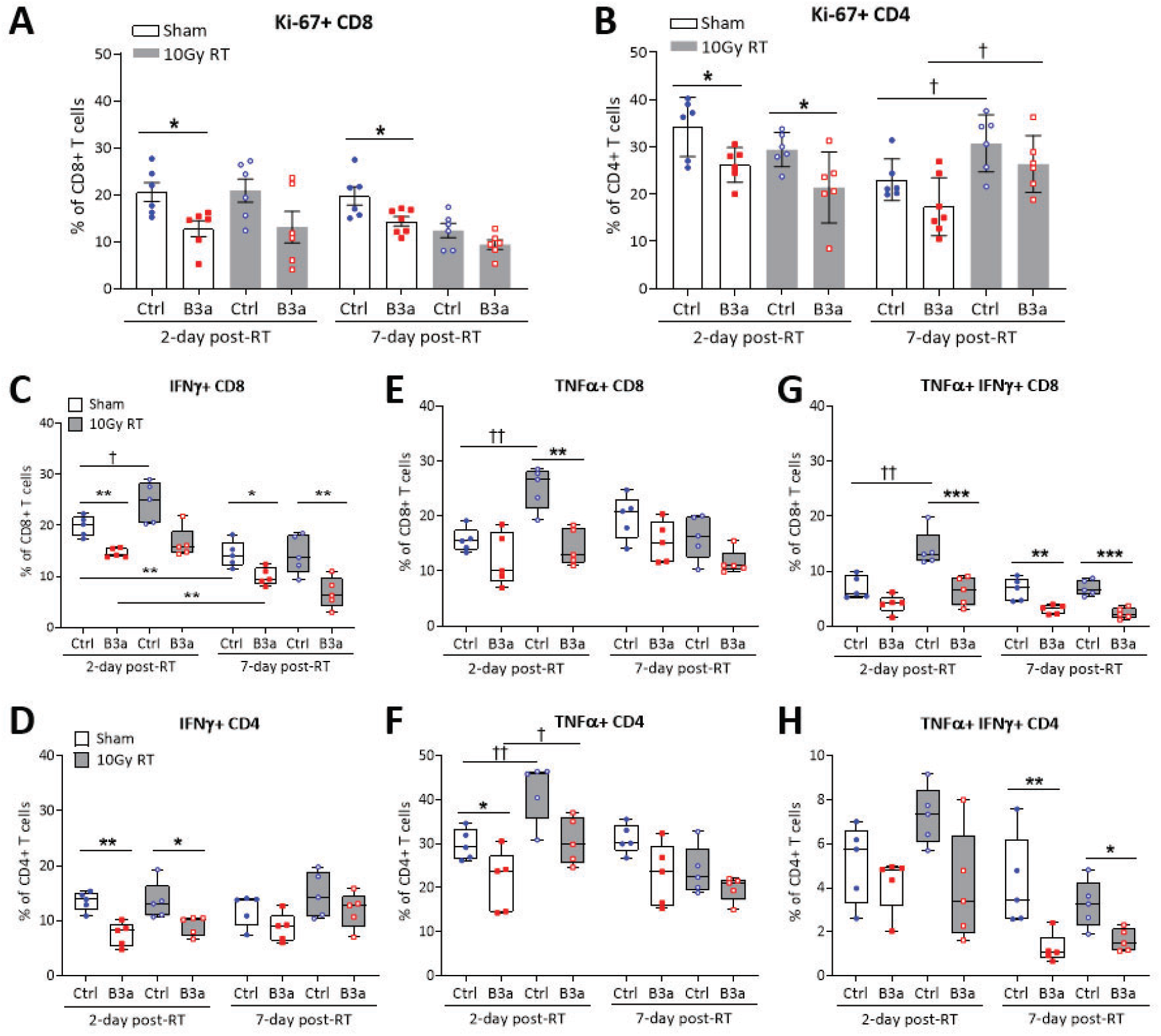
Tumor infiltrating T cells from mSerpinb3a tumors display impaired proliferation and functionally exhausted phenotype. (**A**-**B**) Frequencies of Ki-67+ CD8+ T cells and Ki-67+ CD4+ T cells in total infiltrating CD8 T and CD4 T cell population, respectively. Each dot represents a biologically independent animal. (**C-H**) T cell functionality was assessed by *ex vivo* stimulation with phorbol 12-myristate 13-acetate (PMA)/ionomycin for 5 h and the expression of IFN-ɣ and TNF was examined by intracellular staining. Protein transport inhibitor, brefeldin A was used to block the protein transport processes and cytokine release. Frequencies of (**C**) IFN-ɣ+ CD8 T, (**D**) IFN-ɣ+ CD4 T (**E**) TNFα+ CD8 T, (**F**) TNFα+ CD4 T (**G**) IFN-ɣ and TNFα double positive CD8 T (**H**) IFN-ɣ and TNFα double positive CD4 T were quantified by flow cytometry (n=5 per group). Positive expression was normalized to cells without PMA/ionomycin stimulation (basal levels). Box plot whiskers span minimum and maximum, lines represent median, and each dot represents a biologically independent sample from LL2 tumors. Data are shown as means ± SEM; * indicates comparisons between LL2/Ctrl and LL2/mB3a; ✝ indicates comparisons between sham-treated and RT; \**P* < 0.05, \*\**P* < 0.01, \*\*\**P* < 0.001 using one-way ANOVA with Tukey’s post-test for multiple comparisons.

To examine whether radiation-enhanced T cell response was altered in mSerpinb3-expressing tumors, CD8^+^ and CD4^+^ TILs were stimulated *ex vivo* with PMA/ionomycin and production of tumor necrosis factor alpha (TNFα) and interferon gamma (IFNγ) was evaluated by flow cytometry, as indicative of T cell cytotoxic activity. By stimulating sham-treated CD8^+^ TILs, an average of 20% of CD8^+^ TILs from LL2/Ctrl and 15% from LL2/mB3a tumors showed IFNγ production while the frequencies of IFNγ-producing CD8^+^ TILs were reduced with tumor growth in both groups (Figure 5C). There was no tumor growth-dependent change in IFNγ production in CD4^+^ TILs (Figure 5D), nor in TNFα production in CD4^+^ and CD8^+^ TILs in either tumor background (Figure 5, E and F). Stimulated CD8^+^ TILs taken from irradiated tumors, showed significant enhancement of IFNγ and TNFα production following stimulation in CD8^+^ TILs from LL2/Ctrl tumors at 2-day post-RT but not at 7-day post-RT, whereas radiation-boosted IFNγ and TNFα production was not observed at either time point in CD8^+^ TILs from LL2/mB3a tumors (Figure 5, C and E). Additionally, despite a significant loss of CD4^+^ TILs after radiation in both LL2/Ctrl and LL2/mB3a tumors, their capabilities to produce IFNγ following stimulation remained unchanged, and no radiation-mediated improvement was observed (Figure 5D). In contrast, radiation increased the frequencies of TNFα-producing CD4^+^ TILs following stimulation in both LL2/Ctrl and LL2/mB3a tumor-derived TILs, but similar to TNFα-producing CD8^+^ TILs, the transient improvement was lost by 7-day post RT (Figure 5F). Transient-improved T cell response by radiation was also observed in T cells double-positive for IFNγ and TNFα. Radiation induced a significant increase in the frequencies of IFNγ+TNFα+ CD8 T cells and a non-statistically significant but slight increase in IFNγ+TNFα+ CD4 T cells following stimulation in LL2/Ctrl tumor-derived TILs. However, radiation did not lead to increased frequencies of IFNγ+TNFα+ CD8 or CD4 T cells derived from LL2/mB3a tumor, which showed consistently lower IFNγ and TNFα production than LL2/Ctrl tumor-derived TILs, suggesting a functionally exhausted phenotype (Figure 5, G and H).

### Tumor draining lymph nodes of SERPINB3 tumor-bearing mice have higher Treg and MDSC infiltrates

The TDLN is the site to which tumor antigens first drain and plays a pivotal role in antigen presentation, T cell priming and subsequent tumor infiltration (25). We found that although CD8^+^ T cells were the same in the TDLNs of LL2/mB3a compared to LL2/Ctrl mice (Supplemental Figure 8A), significantly fewer CD8^+^ T cells were seen in LL2/mB3a tumors compared to LL2/Ctrl at 7-day post-RT (Figure 3J), perhaps explained by the high number of suppressive immune cells present in LL2/mB3a tumors discouraging recruitment of CD8^+^ T cells to tumors. Additionally, TDLN of sham-LL2/mB3a tumor bearing mice appeared to have more CD4^+^ T cells at 7 days post-RT, which was then reduced by radiation (Supplemental Figure 8B). However, approximately 20% of these CD4^+^ T cells in both scenarios expressed Treg markers, suggesting a suppressive phenotype of CD4^+^ TDLN T cells in LL2/mB3a tumor bearing mice (Supplemental Figure 8C). MDSCs were not significantly different in the TDLN at early time points, but were increased in mB3a tumors compared to Ctrl tumors at the 7-day timepoint, both in sham- and RT-treated tumors (Supplemental Figure 8D).

### SERPINB3 mediates suppressive chemokine production through STAT signaling activation

Although SERPINB3 has been implicated in pro-inflammatory signaling in pancreatic cancer and Kras mutant tumors (26), the underlying molecular mechanism is unknown. To provide further insight of SERPINB3-mediated suppressive immune response, we focused on understanding the signal activation contributing to suppressive chemokine production in SERIPNB3 cells. We first employed a human phosphorylation pathway profiling array that contained five cancer-associated pathways - MAPK, AKT, JAK/STAT, NF-κB and TGF-β, and identified 14 proteins with upregulated phosphorylation (fold change ≥ 2) and 4 proteins with downregulation (fold change ≤ 0.5) in Caski/B3 compared to Caski/Ctrl cells (Supplemental Figure 9). Among those with increased phosphorylation, signal transducer and activator of transcription (STATs) proteins, including STAT1/2/3/5, showed the highest magnitude of change in phosphorylation (Figure 6A). Increased phosphorylation of STAT1 and STAT3 in Caski/B3 (B3#1, B3#2) relative to Caski parental cells (WT) and Caski/Ctrl (Ctrl) was confirmed by immunoblotting (Figure 6B, untreated). Given that STAT proteins regulate the transcriptional activity of many cytokines and chemokines, we hypothesized that SERPINB3 mediates suppressive chemokine production through promoting STAT signaling activation. Therefore, we inhibited STAT phosphorylation using ruxolitinib, a small-molecule inhibitor approved for clinical treatment by the US Food and Drug Administration for the treatment of myeloproliferative diseases. Caski/WT, Caski/Ctrl, and Caski/B3 were treated with DMSO or ruxolitinib at 1uM and immunoblotting confirmed the inhibition of STAT1 and STAT3 phosphorylation. In untreated and DMSO-treated group, Caski/B3 showed higher levels of STAT1/3 phosphorylation and CXCL1/8 expression compared to Caski/WT and Caski/Ctrl, whereas CXCL1/8 expression was downregulated in Caski/B3 cells treated with ruxolitinib (Figure 6B). Chemokine gene expression and protein secretion were confirmed by qPCR and ELISA, where CXCL1 and CXCL8 were significantly suppressed by ruxolitinib in Caski/B3 (Figure 6, C and E-F), as well as S100A8 and S100A9 (Figure 6, D and G). This may suggest a potential strategy to counteract the suppressive immune response mediated by SERPINB3 in tumor environment.

**Figure 6.**
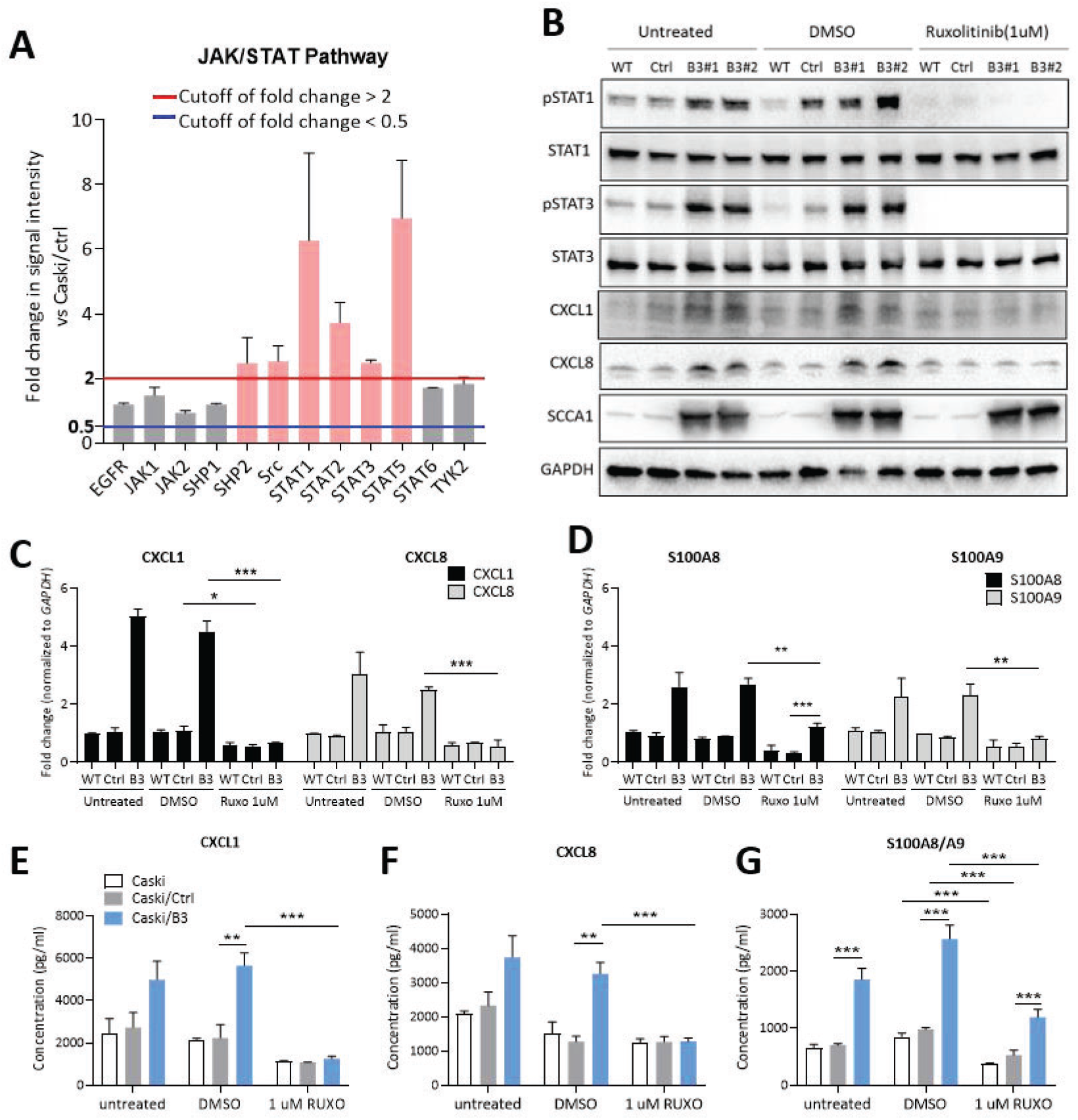
SERPINB3 mediates suppressive chemokine production through STAT signaling activation. (**A**) Activation of JAK/STAT pathway-associated proteins was evaluated using phosphorylation antibody array. Fold changes in phosphorylation were calculated by normalizing the intensity to the expression levels in Caski parental cells and comparing the phosphorylation intensity in Caski/B3 to the levels in Caski/Ctrl cells. Red line indicates fold change ≥ 2 and blue line indicates fold change ≤ 0.5. (**B**) Immunoblotting shows the inhibition of STAT1/3 phosphorylation and CXCL1/8 expression after treating Caski parental cells (WT), Caski/Ctrl (C), and Caski/B3 (B3#1, B3#2) with 1uM Ruxolitinib for 48 h. (**C-D**) Caski/WT, Caski/Ctrl, and Caski/B3 were treated with 1uM Ruxolitinib and the expression CXCL1/8 and S100A8/A9 mRNA was examined by qPCR. Gene expression were normalized to GAPDH and fold changes were calculated by comparing to the expression levels in Caski/WT. (**E-G**) The secretion of (**E**) CXCL1, (**F**), CXCL8, S100A8/A9 after treating with 1uM Ruxolitinib was examined by ELISA. Data in **C-G** are shown as mean ± SEM of n = 4 independent experiments, \**P* < 0.05, \*\**P* < 0.01, \*\*\**P* < 0.001 using one-way ANOVA with Tukey’s post-test for multiple comparisons.

### Elevated serum SCCA and high tumor pSTAT3 is associated with CD11b expression and poor cancer-specific survival after CRT

To evaluate the clinical implication in our findings, tissue microarray containing pre-treatment cervix tumor biopsy specimens obtained from patients with biopsy-proven invasive cervical carcinoma were immunostained for pSTAT3(Tyr705) and myeloid cell marker CD11b. Pretreatment serum SCCA value from 72 cancer patients with an average of 9.16 ng/ml presented a significant cutoff point for cancer specific survival in our patient population. Patients with elevated pretreatment SCCA (⩾9.16 ng/ml) had worse survival than those with low SCCA at the time of diagnosis (Figure 7A), in agreement with our previous study reporting SCCA as a clinical biomarker. The histoscore of pSTAT3 evaluated by immunochemistry was determined through the combined factors of the intensity and percentage of stained cells within the tumor proportion of TMA cores using an attribute cutoff of 100 for high or low pSTAT3 expression (Figure 7B). In high pretreatment SCCA patient cohort (≥9.16 ng/ml), 71% of the population showed high pSTAT3 histoscore as opposed to 41% of high pSTAT3 in low pretreatment SCCA patients (<9.16 ng/ml) (Figure 7C). Althoug hp STAT3 was not an independent prognostic factor for survival in our cohort, the patients with elevated serum SCCA, along with high pSTAT3 were associated with increased CD11b expression (Figure 7D). In contrast, the majority of patients with pretreatment SCCA <9.16 ng/ml had low pSTAT3 histoscore and was correlated with low CD11b expression. This cohort had the highest cancer-specific survival (Figure 6A). Overall, SCCA is a strong clinical biomarker and when combining with pSTAT3 expression, it may indicate an unfavorable tumor microenvironment and provide an opportunity for selection of patients for anti-STAT and/or anti-SERPINB3 directed therapies.

**Figure 7.**
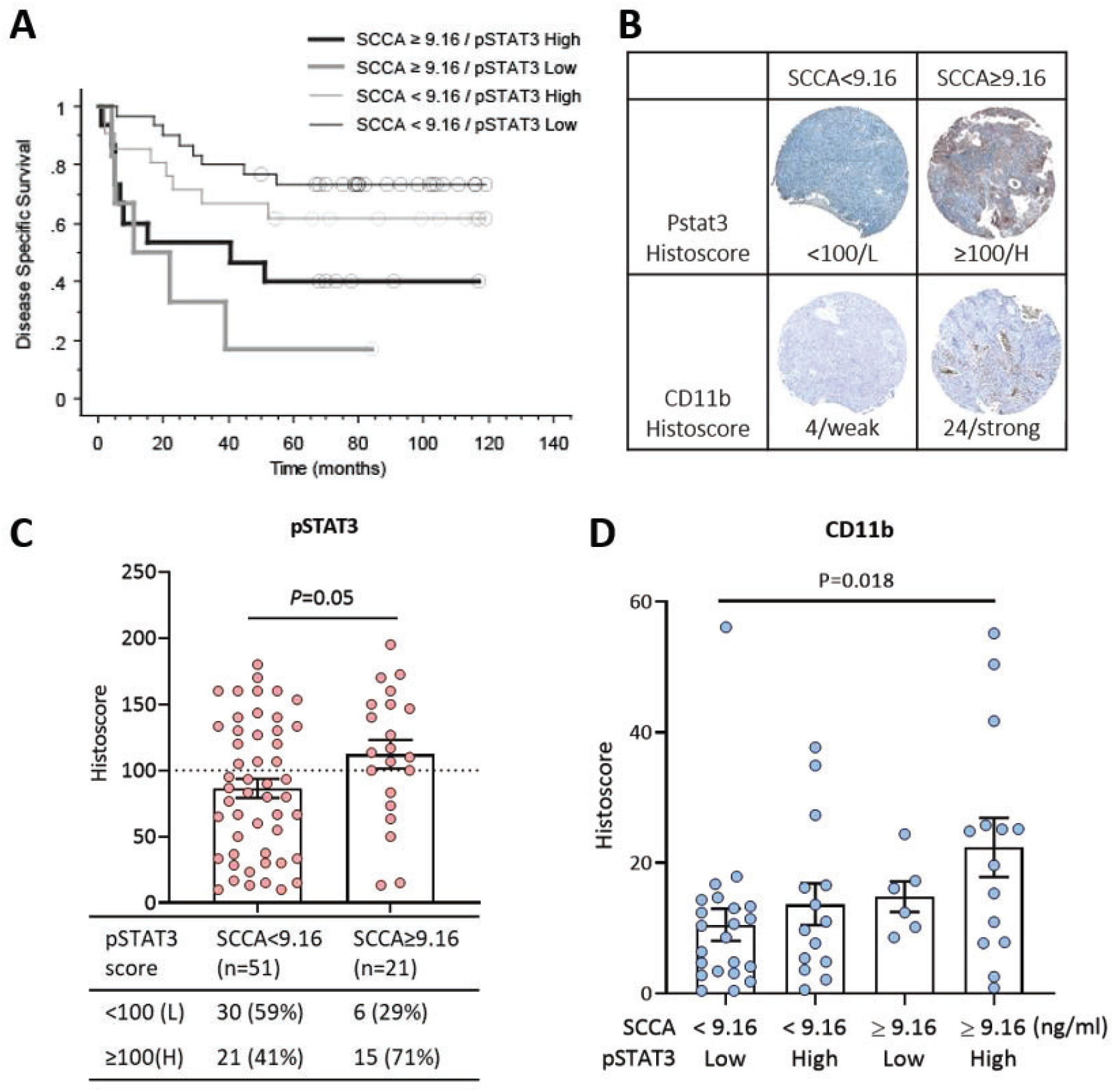
Elevated serum SCCA and high tumor pSTAT3 is associated with CD11b expression and poor cancer-specific survival after CRT. (**A**) Kaplan-Meier plot show overall survival in patients with serum SCCA1 <9.16 ng/ml with either pSTAT3 histoscore <100 (n=30) or ≥100 (n=21), compared to patients with serum SCCA1 ≥9.16 ng/ml with either pSTAT3 histoscore <100 (n=6) or ≥100 (n=15). The average pretreatment serum SCCA value of 9.16 ng/ml from 72 cancer patients was used as a cutoff. (**B**) Representative images of pSTAT3 and CD11b staining from patients with SCCA1 < or ≥9.16 ng/ml **(C**) Phosphorylated STAT3 staining score (histoscore) in patients with serum SCCA1 <9.16 ng/ml versus SCCA1 ≥9.16 ng/ml. (**D**) Myeloid cell marker, CD11b staining score in patients with serum SCCA1 < or ≥9.16 ng/ml with pSTAT3 histoscore <100 (low) or ≥100 (high). Each dot represents an individual patient. Data are shown as means ± SEM, using unpaired two-tailed Student’s t-test.

## Discussion

In this study, we revealed that SERPINB3 modulated tumor microenvironment towards a pro-tumor phenotype by regulating suppressive chemokine production, to facilitate tumor growth and impede the success of RT. Genetically inducing SERPINB3 in human cancer cells led to increased production of CXCL1/8 and S100A8/A9, which promoted myeloid cell migration. These chemokines were increased in the tumors of both human SERPINB3-expressing Caski xenograft and murine Serpinb3a-expressing LL2 syngeneic mouse models, resulting in substantial numbers of infiltrating M-MDSCs, PMN-MDSCs, and M2 macrophages. Radiation-induced T cell responses were compromised by suppressive microenvironment in SERPINB3 tumors. It is worth noting that the correlation between SERPINB3 and CXCL1/8, S100A8/A9 was conserved across several cancers known to have high SERPINB3 expression and often associated with poor treatment outcomes, suggesting wide application of our study to a variety of tumors with SERPINB3 expression. We also revealed a regulatory mechanism of STAT-dependent chemokine expression by SERPINB3; therefore, targeting STAT signaling might prevent SERPINB3-mediated immunosuppression.

CXCL1/8 are potent mediators of immune cell chemotaxis via chemokine receptor CXCR2, commonly expressed on myeloid progenitor cells, neutrophils, monocytes and macrophages (27). S100A8 and S100A9 are well-established immunosuppressive factors in tumors, known to attract MDSC infiltration, promote suppressive macrophage differentiation, and stimulate immunosuppressive activity (28-30). Our findings that SERPINB3 modulates the crosstalk between immune and cancer cells via secretion of CXCL1/8 and S100A8/A9 implicates this protease inhibitor member of the SERPIN superfamily in a key tumor strategy to evade the anti-tumor immune response and resist therapies such as radiation. Our findings correspond to previous studies of SERPINB3 in atopic dermatitis and psoriasis, whereby downregulation of SERPINB3 in keratinocytes was associated with reduced expression of CXCL1, 5, 8 (31), and S100A8 (18). Catanzaro and colleagues showed that SERPINB3 was a downstream mediator of mutant Ras-induced tumorigenesis and knocking down SERPINB3 expression led to decreased IL-6, CXCL1, CXCL8 production suppressing tumorigenesis (26). However, the downstream immune consequences were not further examined in these studies. Moreover, high SERPINB3 is associated with lymph node metastasis in patients with cervical cancer (32, 33) and esophageal squamous cell carcinoma (34), but the underlying causes are not fully understood. Given that S100A8/A9 is a crucial player in orchestrating immunosuppression in distant organs and establishing a pre-metastatic niche by priming tissues for tumor cell deposition (35, 36), high levels of CXCL1 and S100A8/A9 in SERPINB3-tumor bearing mice, along with increased accumulation of MDSCs in TDLNs provided a possible mechanism for high metastatic tendency of SERPINB3 tumors.

Although in vivo experiments were designed to measure the short-term effects of radiation on tumor immune infiltrates in mB3a versus Ctrl tumors, we did see some evidence that radiation-induced tumor growth delay was blunted in the mB3a tumors. This is consistent with our previous experiments powered and designed to detect this difference that SERPINB3 expression inhibits tumor growth delay (12). In our syngeneic models, we did not observe radiation-associated T cell expansion, an important radiation-induced response to enhance antitumor immunity (37-39). Less Ki-67+ CD8+ TILs in mB3a tumors compared to Ctrl was mainly attributed to the pre-existing TME, and independent of radiation effect. The expression of Ki-67 in CD4+ TILs was increased at 7-day post-RT; however, given that Treg cells are a subpopulation of CD4+ TILs, increased Ki -67 expression might be associated with the expansion of Treg cells which have a more radioresistant properties than other T lymphocytes (24). Ratio of CD8 to Treg served as a predictive factor for clinical response in many cancer patients; for instance, high CD8/Treg ratio correlated with favorable outcome in cervical cancer treated with neoadjuvant chemotherapy (40), improved survival in oral squamous cell carcinoma (41), and radiosensitivity in squamous cell carcinoma of the larynx (42). In both mB3a and Ctrl tumors, CD8+ TIL percentage increased at 7-day post-RT but this increase in mB3a tumors failed to result in higher CD8/Treg ratio, as shown in Ctrl tumors, suggesting a higher degree of Treg cell expansion in mB3a tumors induced by radiation.

In addition to higher levels of suppressive myeloid lineage cells, we also showed that SERPINB3 tumors harbored functionally exhausted T cells that failed to demonstrate RT-induce T cell responses. Although inhibitory T-cell receptor signaling was not investigated in this study, some suggest a potential connection between SERPINB3 and PD-L1 expression. In HPV-negative head and neck squamous cell carcinoma patients from TCGA database, high SERPINB3 expression corresponded to increased PD-L1 and PD-L2 (43). Similarly, genome-level and IHC showed upregulated PD-L1 in SERPINB3-high ovarian and esophageal tumors (44). The mechanism of how SERPINB3 may result in inhibition of T cell proliferation, function, and exhaustion phenotype is unknown, and is a focus of ongoing research. This current study revealed the increased infiltration of M-/PMN-MDSCs, TAMs and M2 macrophages in SERPINB3-high tumor, and that tumor-directed irradiation further augmented the immunosuppression by inducing CXCL1, S100A8/A9 production and M-/PMN-MDSCs infiltration. Future therapies may involve targeting of these chemokines, immune cell subsets, or even upstream molecules like SERPINB3 for maximal tumor-specific effects.

Furthermore, inhibiting STAT signaling may be one promising approach to target SERPINB3-modulated immunosuppression. Despite evidence that the JAK inhibitor ruxolitinib successfully triggered tumor regression, increased drug sensitivity, prevented angiogenesis in many preclinical models, there may be several explanations for why clinical trails in solid tumors to date have been largely unsuccessful (45). Few trials have combined ruxolitinib with radiation. One study showed a promising result from phase I study that glioma patients received the treatment of ruxolitinib with radiation and temozolomide had significantly better overall survival than those that received ruxolitinib and radiation (46). The consideration of using STAT inhibitor in combination with radiation therapy was also found in an ongoing clinical trial (NCT01904123) using STAT3 inhibitor, WP1066. Preclinical murine models showed that STAT3 inhibitor and irradiation reprogrammed immunosuppressive glioma microenvironment by improving dendritic cell maturation and interactions with T cells, resulting in enhanced survival compared to either treatment alone (47). In addition, inhibiting STAT3 activity in myeloid cells improved RT-induced anti-tumor immunity in murine xenograft models of head and neck cancers (48). Whether synergizing with radiation to evoke antitumor immunity was critical for improving the efficacy or it was cancer type-specific treatment response requires further investigation. Additionally, as STAT transcriptional activity is also involved in T-cell function (49), and other facets of the immune response, direct inhibition of this pathway may unintentionally tip the immune axis back in favor of the tumor. Thus, targeting upstream, tumor-specific signals such as SERPINB3, may be more effective in the clinical setting.

In conclusion, SERPINB3 modulated suppressive immune milieu by enhancing CXCL1/8 and S100A8/A9 expression through STAT signaling to promote the infiltration of MDSCs, M2 macrophages, Treg cells, and simultaneously disrupt CD8+ T cell function. As a result, the beneficial immunological response induced by radiation was muted in SERPINB3 tumors. Future study on targeting STAT signaling and our ongoing endeavor to develop SERPINB3-targeting drugs are warranted to improve RT-induced antitumor immunity.

## Methods

### Cell lines and plasmids

Cell lines were purchased from the ATCC. Caski, SW756, C33a were cultured in Iscove’s Modified Dulbecco’s Medium (IMDM) and Lewis lung carcinoma (LL2/LLC) were cultured in Dulbecco’s modified Eagle’s medium (DMEM), supplemented with 10% fetal bovine serum (FBS), and 100 U/mL penicillin-streptomycin. Generation of SERPINB3 CRISPR-Cas9 knockout cells was described previously (12). SERPINB3 stable expression cells were generated using pULTRA lentiviral vector (Addgene #24129) containing human SERPINB3 or pLV-C-GFPSpark vector (Sino Biological LVCV-35) containing mouse Serpinb3a GFP-tagged fusion proteins. SERPINB3 knockdown cells were transduced with scramble shRNA (Addgene #1684) or SERPINB3 shRNA (Sigma Mission shRNA, TRCN0000052400). Genetically modified cells were generated through a lentivirus system by transfection of human 293T packaging cells. All cell lines were grown in monolayer at 37°C with 5% CO2 and periodically tested for Mycoplasma contamination

### RNA sequencing and TCGA data analysis

RNA sequencing (RNAseq) was performed on pre-treatment tumor biopsies obtained from patients enrolled on a prospective tumor banking study with written informed consent (Washington University IRB No. 201105374). RNAseq processing and normalization has been reported previously (50) and data are available on GeneExpression Omnibus (GEO): accession number: GSE151666. Immune cell population and enrichment score were analyzed using xCell analysis, a gene signature-based method to estimate cell composition in bulk transcriptomic data (22). Heatmaps were generated using GraphPad Prism base on the average scores for each immune cell subtypes in our predefined patient groups. The Cancer Genome Atlas(TCGA) RNAseq data were obtained through cBioPortal (http://www.cbioportal.org/) www.cbioportal.org/). Spearman’s correlation coefficient was used to test the significance of the correlation with an adjusted P<0.05.

### Serum SCCA and tissue microarray immunohistochemistry

Pretreatment serum SCCA was evaluated by ARUP National Reference Laboratory (Salt Lake City, UT, USA) using ELISA and tissue microarray was generated from untreated tumor specimens, as described previously (16). TMA sections were sent to HistoWiz Inc for IHC staining for CD11b (1:100, Abcam ab224800) and IHC for pSTAT3 (1:200, Sigma SAB4300033) was performed by Washington University AMP Core Labs. QuPath V0.3.2 software was used for automated analysis using surface and cytoplasmic staining to determine percent cells positive for CD11b. The staining scores for pSTAT3 were evaluated by the pathologist and calculated by (% of positive stained tumor cells x staining intensity ranged from 0-3). Values from at least two cores from each patient were considered valid and an average score was taken.

### PBMC isolation and transwell assay

Fresh primary PBMCs were obtained from patients planned to undergo radiation therapy with brachytherapy for cervical cancer and had enrolled on a prospective biospecimen banking protocol (Institutional Review Board Protocol #201105374). A total volume of 10-20 ml of fresh blood was collected in EDTA separator tubes and peripheral blood mononuclear cells (PBMC) were immediately isolated using Lymphoprep and SepMate-50 (Stemcell Technologies) centrifugation tubes, according to the manufacture’s instruction. PBMCs were cultured in RPMI 1640 supplemented with 10% heat-inactive FBS and 100 U/mL penicillin-streptomycin. Transwell assay was performed using 8-μm Transwells (Falcon). Supernatants were collected from cells cultured in complete growth media for 48 h and loaded in the lower chamber of the transwell. PBMCs were loaded to the upper transwell for a 4 h migration period. Migrated cells were phenotyped by flow cytometry.

### Flow cytometry and data analysis

Single cell suspensions were blocked with either Human TruStain FcX Solution (422301, Biolegend) or mouse TruStain FcX PLUS (anti-mouse CD16/32) Antibody (S17011E, bioLegend) to avoid nonspecific Fc receptor binding and stained with LIVE/DEAD Fixable Dead Cell Stain Kit (MACS) to exclude dead cells. For surface staining, cells were incubated with the appropriate antibodies for 30 min at 4 °C. Intracellular cytokine and nuclear staining was performed after surface staining using Cyto-Fast Fix/Perm Buffer Set and True-Nuclear Transcription Factor Buffer Set, respectively (BioLegend). Stained cells were analyzed using MACSQuant Analyzer 10 Flow Cytometer. Antibody information is shown in Supplemental Table S2. Data analysis including viSNE and FlowJo plugin FlowSOM were performed on FlowJo v.10 (TreeStar). A range of 20,000 to 80,000 live cells was acquired and individual flow cytometry data from each group were combined into a single data file for generating viSNE. Color-coded subpopulations were gated by pre-defined markers for each immune cell types and overlaid to the viSNE plots for total live cells from tumors. All flow cytometry gating plots, histograms, and statistics were generated on FlowJo and exported to GraphPad Prism v.8 for graph visualization.

### Mouse tumor models and radiation

Female athymic nude mice (Charles River) between 6 and 7 weeks old were injected subcutaneously with 5×10^6^ Caski/Ctrl or Caski/B3 cells suspended in serum-free IMDM and 50% Matrigel Basement Membrane Matrix (Corning) to a final volume of 100μl on their flank. Female C57/BL6 mice (Charles River) aged 7-8 weeks were injected subcutaneously with 5 × 10^5^ LL2/Ctrl or LL2/mB3a cells suspended in 50ul of serum free DMEM into the right flank. Mice were sacrificed on the day indicated or randomized to receive sham or 10Gy RT. Tumor volume was measured twice weekly and calculated by (length × width^2^)/2. Mouse tumors were irradiated using the Xstrahl Small Animal Radiation Research Platform (SARRP) 200 (Xstrahl Life Sciences). Animal work was approved by the Washington University Institute Institutional Animal Care and Use Committee (Protocol #20-0470). To characterize immune infiltrates, tumors were manually dissected and digested with 1 mg/ml Collagenase, 0.5 mg/ml hyaluronidase, and 10 mg/mL DNase I type IV (Sigma), and transferred to a tissue disaggregator Medicon (Becton Dickinson) using CTSV Medimachine II (Becton Dickinson) for cell dissociation and tissue homogenate was filtered through a 100 μm strainer. Lymph nodes were dissociated by being pressed through a 70 μm filter. Flow cytometry was used for immune cell characterization.

### Ex vivo T cell stimulation

Single-cell suspensions from tumors or CD3 T cells isolated using MojoSort Mouse CD3 T Cell Isolation Kit (Biolegend) following the manufacture’s protocols, were stimulated with 500x Cell Activation Cocktail containing 40.5 μM phorbol-12-myristate 13-acetate (PMA) and 669.3 μM ionomycin (Biolegend) in the presence of 5 μg/mL BFA (Biolegend) in RPMI media supplemented with 10% HI FBS. Cells were stimulated for 5 h and stained with surface/intracellular markers for flow cytometry analysis.

### RNA extraction and qPCR

RNA was isolated using GenElute Mammalian Total RNA Miniprep Kit (Sigma) and reverse transcribed to cDNA using High Capacity cDNA Reverse Transcription Kit(Invitrogen). Quantitative polymerase chain reaction (qPCR) was performed using PowerUp SYBR green PCR Master Mix (Applied Biosystems) and the Applied Biosystems 7900 Fast real-time PCR system and software. Each sample was performed in triplicate, gene expression levels were normalized to GAPDH and fold changes were calculated using the ΔΔCt method. Sequences of primers are detailed in Supplemental Table 3.

### Enzyme-linked immunosorbent assay (ELISA)

Cell culture supernatant was collected at 48 h after fresh media was added to the adherent cells in monolayer. Cell lysates were prepared using NP-40 buffer (Alfa Aesar). Quantification of human/mouse chemokines in tissue culture supernatants and tissue homogenates was performed using a commercially available human CXCL1/GRO alpha, human IL-8/CXCL8, human S100A8/S100A9 Heterodimer, mouse CXCL1/KC, and mouse S100A8/S100A9 Heterodimer DuoSet ELISA kit from R&D Systems. Mouse GRO gamma ELISA Kit was obtained from Abcam. Chemokine concentration in samples was determined by interpolation from a standard curve.

### Phosphorylation protein array

Human Phosphorylation Pathway Profiling Array C55 consisted of the detection of 55 phosphorylated proteins (RayBiotech). Same amount of protein from each sample was used for screening and assays were performed according to the manufacturer’s instruction. Array blots were scanned with Bio-Rad ChemiDoc MP imaging system and images were processed using Protein Array Analyzer plug-in (http://image.bio.methods.free.fr/ImageJ/?Protein-Array-Analyzer-for-ImageJ.html) of the ImageJ program.

### Immunoblotting

Cells were treatedwithruxolitinib (Apexbio Technology) and lysed with RIPA buffer (Cell signaling) supplemented with proteinase/phosphatase inhibitors (Thermo). Proteins concentration was determined using BCA (Thermo) and proteins were electrophoresed on 4–20% gradient gels (Bio-Rad), transferred to PVDF blot using the Trans-Blot TurboTransfer system (Bio-Rad), and incubated with antibodies shown in Supplemental Table 2. Chemiluminescence was detected by using ECL reagent (Cytiva) and visualized using the Bio-Rad ChemiDoc MP imaging system and Image Labsoftware (Bio-Rad).

### Statistics and data availability

Statistical analyses was performed using GraphPad Prism version 8 and all values are reported as mean ± SEM. Two-tailed unpaired t test was used for two groups comparisons and one-way or two-way ANOVA was used for multiple comparisons. P values of less than 0.05 were considered statistically significant (*P < 0.05; **P < 0.01; ***P < 0.001). All the data acquired in the study are available from the corresponding author.

### Data Availability

The data generated in this study are available within the article and its supplementary data files. All raw data are available upon request from the corresponding author. RNA sequencing data are available on GeneExpression Omnibus (GEO): accession number: GSE151666. RNA sequencing from The Cancer Genome Atlas (TCGA) consortium was obtained through cBioPortal (http://www.cbioportal.org/).

## Author contributions

LC and SM conceptualized the study. LC designed and conducted experiments, acquired data, performed formal analysis, and wrote the manuscript. VS, SW, and RF conducted experiments and acquired data. FR, KJ, JZ, and PC contributed to RANseq data curation and analysis. LS interpreted and analyzed histology data. CJL, CS, PWG, and JKS provided resources. SM supervised the study, acquired funding, and wrote the manuscript. All authors reviewed and edited the manuscript.

## Acknowledgments

This work was supported by NIH grant K08CA237822, American Society for Clinical Oncology (ASCO) Career Development Award #13170, American Cancer Society Institution Research Grant IRG1815861, and Washington University Radiation Oncology Seed Grant. We would like to acknowledge Washington University Brachytherapy Nurses and Staff for assisting in collection of biospecimens. We would also like to thank Dr. Cedric Mpoyand Dr. Buck Rogers of the Small Animal Radiation Research Platform Core for their assistance with animal experiments, Dr. Dinesh Thotala for gifting the mouse Lewis lung carcinoma cell line and Lena Zein for administrative assistance.

